# Atg15 is a vacuolar phospholipase that disintegrates organelle membranes

**DOI:** 10.1101/2023.05.31.543165

**Authors:** Yasunori Watanabe, Yurina Iwasaki, Kyoka Sasaki, Kuninori Suzuki

**Author notes:** To whom correspondence should be addressed: Yasunori Watanabe, Faculty of Science, Yamagata University, 1-4-12 Kojirakawamachi, Yamagata, Yamagata 990-8560, Japan. Kuninori Suzuki, Department of Integrated Biosciences, Graduate School of Frontier Sciences, The University of Tokyo, FSB-101, 5-1-5 Kashiwanoha, Kashiwa, Chiba 277-8562, Japan.

## Abstract

Atg15 (autophagy-related 15) is a vacuolar phospholipase essential for the degradation of cytoplasm-to-vacuole targeting (Cvt) bodies and autophagic bodies, hereinafter referred to as intravacuolar autophagic compartments (IACs), but it remains unknown if Atg15 directly disrupts IAC membranes. Here we show that the recombinant *Chaetomium thermophilum* Atg15 lipase domain (CtAtg15(73–475)) possesses phospholipase activity that in acidic environments digests phospholipid to generate lysophospholipid and free fatty acid. The activity of CtAtg15(73–475) was markedly elevated by limited digestion with proteinase K. Proteinase K–treated CtAtg15(73–475) was detected as three fragments, with cleavage between S159 and V160, and between F209 and N210. We inserted the human rhinovirus 3C protease recognition sequence into the two sites and found that cleavage between S159 and V160 was important to activate CtAtg15(73–475). We confirmed that CtAtg15 compensated for the absence of *ATG15* in *Saccharomyces cerevisiae* cells, indicating that CtAtg15 could disintegrate *S. cerevisiae* IAC *in vivo*. Further, both mitochondria and IAC were disintegrated by CtAtg15 in *S. cerevisiae*. This study suggests that Atg15 can degrade any organelle membrane, indicating that Atg15 plays a role in disrupting organelle membranes delivered to vacuoles by autophagy.

## Introduction

In *Saccharomyces cerevisiae,* a type of selective autophagy called cytoplasm-to-vacuole targeting (Cvt)^1^ is active during growth, and macroautophagy (hereinafter referred to simply as ‘autophagy’) is induced by starvation or rapamycin treatment^2,3^. In both types of autophagy, the vacuolar hydrolase aminopeptidase I (Ape1) is selectively transported to vacuoles^4^. Ape1 is synthesised in a precursor form (prApe1) that assembles into the Ape1 complex in the cytoplasm^5–7^. The Ape1 complex is transported to vacuoles by the Cvt pathway under nutrient-rich conditions, but by autophagy under starvation conditions^4^. In both pathways, two double-membrane compartments, the Cvt vesicles and autophagosomes, are responsible for Ape1 transport *via* the Cvt pathway and autophagy, respectively^4^. The outer membranes of Cvt vesicles or autophagosomes fuse with the vacuolar membrane, and then Cvt bodies or autophagic bodies are released into the vacuole. After disintegration of the limiting membranes of Cvt bodies or autophagic bodies, the propeptide in prApe1 is cleaved by vacuolar proteases; consequently, prApe1 is converted to mature Ape1 (mApe1)^8^. The decrease in the molecular mass of prApe1 by cleavage of the propeptide, as detected by immunoblot analysis, can be used as an indicator of the degradation of Cvt bodies and autophagic bodies. Autophagy is a bulk degradation system that is widely conserved among eukaryotes, from yeast to mammals^2^. It enables cells to survive severe environmental conditions, for instance nutrient limitation, by recycling cellular building blocks such as amino acids and nucleic acids^9–11^.

To degrade the contents enclosed by Cvt bodies and autophagic bodies, the limiting membranes of Cvt bodies and autophagic bodies must be disintegrated in order to allow access by vacuolar hydrolases. Atg15/Aut5/Cvt17, a glycosylated integral membrane protein that is essential for the disintegration of Cvt bodies and autophagic bodies, is the only vacuolar lipase in *S. cerevisiae*^12,13^. Atg15 has a transmembrane domain (TMD) at its N-terminus and a lipase domain at the C-terminus^12^. The S332 residue in the C-terminal domain is thought to be the active center for Atg15 lipase activity^12–14^. Atg15 is localised to the endoplasmic reticulum (ER) and is transported to vacuoles *via* the multivesicular body pathway^12^, but its extreme N-terminal cytoplasmic region (residues 2–12) is not required for vacuolar targeting^15^. Recently, our group showed that the N-terminal TMD is important and sufficient for the vacuolar localisation of Atg15, and targeting of the C-terminal lipase domain to the vacuolar lumen is also important for the degradation of Cvt bodies and autophagic bodies^16^.

Pep4 and Prb1, two vacuolar proteases of *S. cerevisiae*^17–19^, are also required for disintegration of Cvt bodies and autophagic bodies^20^. The function of Pep4 in vacuoles is to activate other vacuolar hydrolases, such as Pho8, Prb1, and Prc1, and to degrade cellular components^17^. Although Pep4 and Prb1 are unlikely to have lipase activity, Cvt bodies and autophagic bodies remain intact in *pep4*Δ and *prb1*Δ cells. A longstanding question in the field is how vacuolar proteases contribute to the disintegration of intravacuolar autophagic compartments (IAC), including Cvt bodies and autophagic bodies.

In this study, we showed that the lipase domain of *Chaetomium thermophilum* Atg15 (CtAtg15(73–475)) possessed phospholipase activity and was activated by cleavage between S159 and V160 by proteinase K. *In vitro* experiments using recombinant CtAtg15 and the membrane fraction of *S. cerevisiae* cells clearly showed that CtAtg15 could disintegrate mitochondrial membranes other than IACs. We conclude that activated Atg15 can disintegrate various kinds of organelles enclosed by phospholipid membranes, suggesting that Atg15 contributes to the disruption of any organelle membranes delivered to vacuoles.

## Results

### The C-terminal lipase domain in Atg15 exhibits phospholipase activity *in vitro*

Recently, our group reported that the transport of the C-terminal lipase domain (residues 50–466) of *S. cerevisiae* Atg15 (ScAtg15) to the vacuolar lumen is sufficient for its lipase activity^16^. The AlphaFold2-predicted structure of ScAtg15 indicated that residues 50–466 were structured^21,22^, supporting the theory that this domain acts as a lipase. In addition to ScAtg15, we predicted the structures of *Kluyveromyces marxianus* Atg15 (KmAtg15) and *C. thermophilum* Atg15 (CtAtg15) with ColabFold^23^ (Fig. S1A–C). Based upon the predicted structures, we prepared the lipase domains of ScAtg15(50– 471), KmAtg15(54–485), and CtAtg15(73–475) as recombinant proteins.

The phospholipase activity of ScAtg15 can be estimated using NBD-PE as a substrate^14^. We examined the activity of the recombinant Atg15 proteins and found that ScAtg15(50–471) and KmAtg15(54–485) showed minimal lipase activity, whereas that of CtAtg15(73–475) was much higher (Fig. 1A). Since CtAtg15 showed higher activity than ScAtg15 or KmAtg15, we decided to use CtAtg15 for *in vitro* analysis. Two kinds of spots were detected by the reaction of CtAtg15 (products 1 and 2 in Fig. 1A). We analysed the spots by mass spectrometry and confirmed that products 1 and 2 corresponded to NBD-free fatty acid (FFA) and NBD-lysoPE, respectively (Fig. 1B–C). This result suggests that CtAtg15 possesses phospholipase B activity. To characterise the pH dependence of CtAtg15, we next examined the activity of CtAtg15 under acidic to neutral pH conditions (Fig. S2A). The activity of CtAtg15 was maximal at pH 4.0 and almost absent at pH 6.0. In ScAtg15, the S332 residue is predicted to be an active center, and the mutation of S332 to alanine abolishes the activity of ScAtg15^12,13,16^. We substituted the corresponding 330 serine residue in CtAtg15 with alanine to generate the CtAtg15(S330A) mutant, and examined the activity of the recombinant CtAtg15(S330A) (Fig. S1D). CtAtg15(S330A) could not hydrolyse NBD-PE (Fig. 1D), indicating that S330 is an active center of CtAtg15 and that CtAtg15 catalyses NBD-PE as a phospholipase. Next, we examined whether CtAtg15 catalysed two other phospholipids, namely phosphatidylserine (PS) and phosphatidylcholine (PC), and found that CtAtg15 could degrade both (Fig. 1E–F). Atg15 does not appear to exhibit substrate specificity for any particular phospholipid. Finally, we showed that CtAtg15 degraded NBD-PE in liposomes (Fig. S2B), suggesting that CtAtg15 can recognise membrane phospholipids.

**Figure 1.**
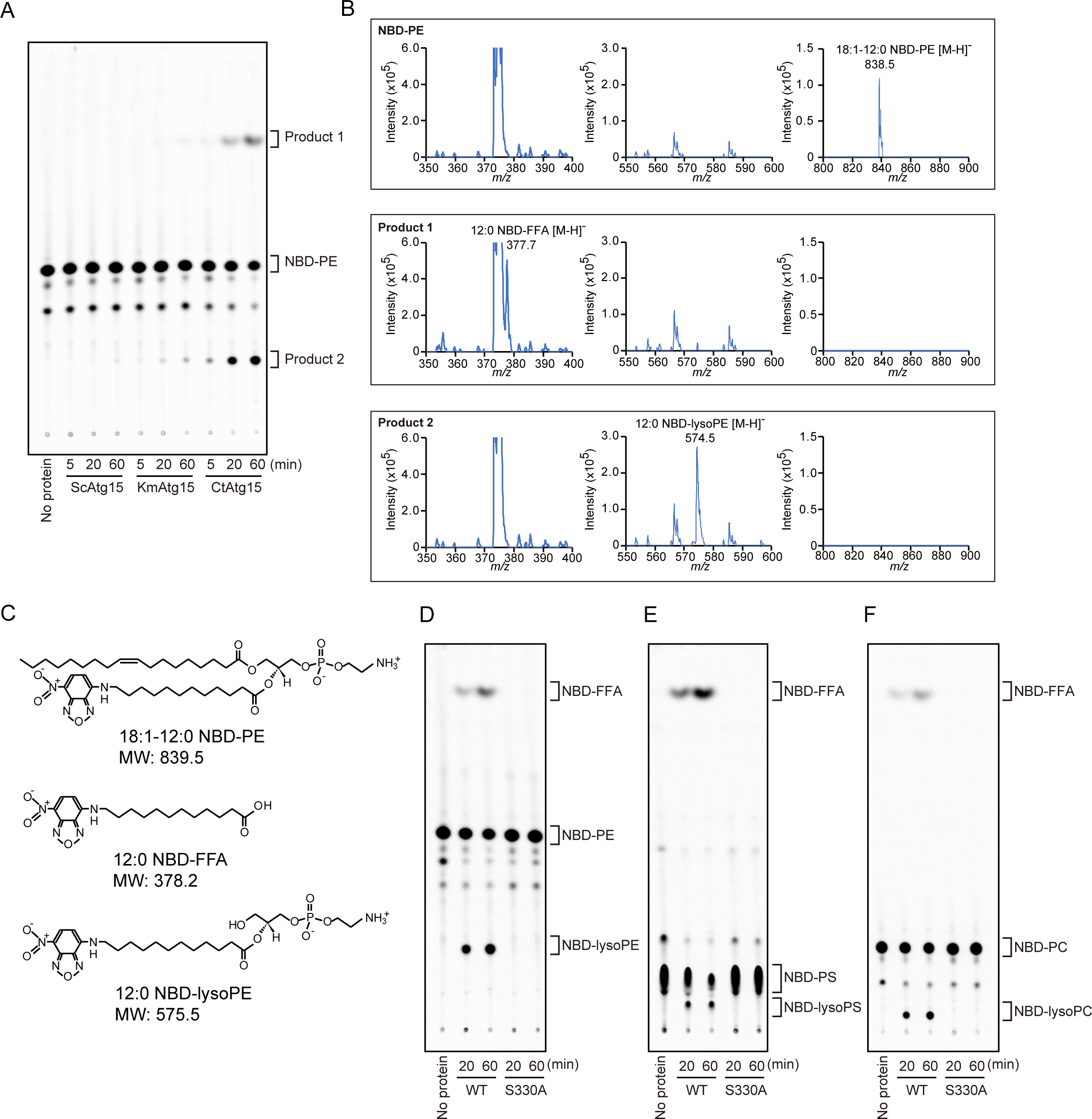
Phospholipase activity of CtAtg15. (A) NBD-PE and purified ScAtg15(50–471), KmAtg15(54–485) and CtAtg15(73–475) were incubated at 30°C for the indicated times. Phospholipids were extracted and analysed by TLC. (B) After TLC separation of products 1 and 2 produced by CtAtg15 (shown in A), products 1 and 2 and NBD-PE were scraped from the TLC plate and extracted. Extracted NBD-PE (upper panel) and products 1 and 2 (middle and lower panels, respectively) were analysed by mass spectrometry using negative ion mode. NBD-PE, NBD-FFA and NBD-lysoPE were detected as the singly charged ions indicated in the charts. (C) Chemical structures of 18:1-12:0 NBD-PE, 12:0 NBD-FFA and 12:0 NBD-lysoPE. (D–F) Substrate specificity of CtAtg15. (D) NBD-PE, (E) NBD-PS and (F) NBD-PC were incubated with the purified wild-type (WT) or the S330A mutant of CtAtg15(73–475) at 30°C for the indicated times. Phospholipids were extracted and analysed by TLC.

### The phospholipase activity of recombinant Atg15 is increased by limited proteolysis

In *S. cerevisiae*, the transport of the C-terminal lipase domain to the vacuolar lumen is sufficient for its lipase activity, but little activity is detected without Pep4^16^. This suggests that cleavage by a protease in the lipase domain is needed to activate Atg15. Previous studies have shown that vacuolar carboxypeptidase Y can be activated by proteolysis via *S. cerevisiae* Pep4^24^ or proteinase K^25^. Since proteinase K activates carboxypeptidase Y more effectively than Pep4^25^, we decided to use the former in our experiments. We compared the lipase activity of ScAtg15(50–471), KmAtg15(54–485) and CtAtg15(73–475) with or without proteinase K treatment. As expected, the activity of all Atg15 species was significantly elevated by treatment with proteinase K (Fig. 2A– B). This result supports our hypothesis that processing of the Atg15 lipase domain is important for its activation.

**Figure 2.**
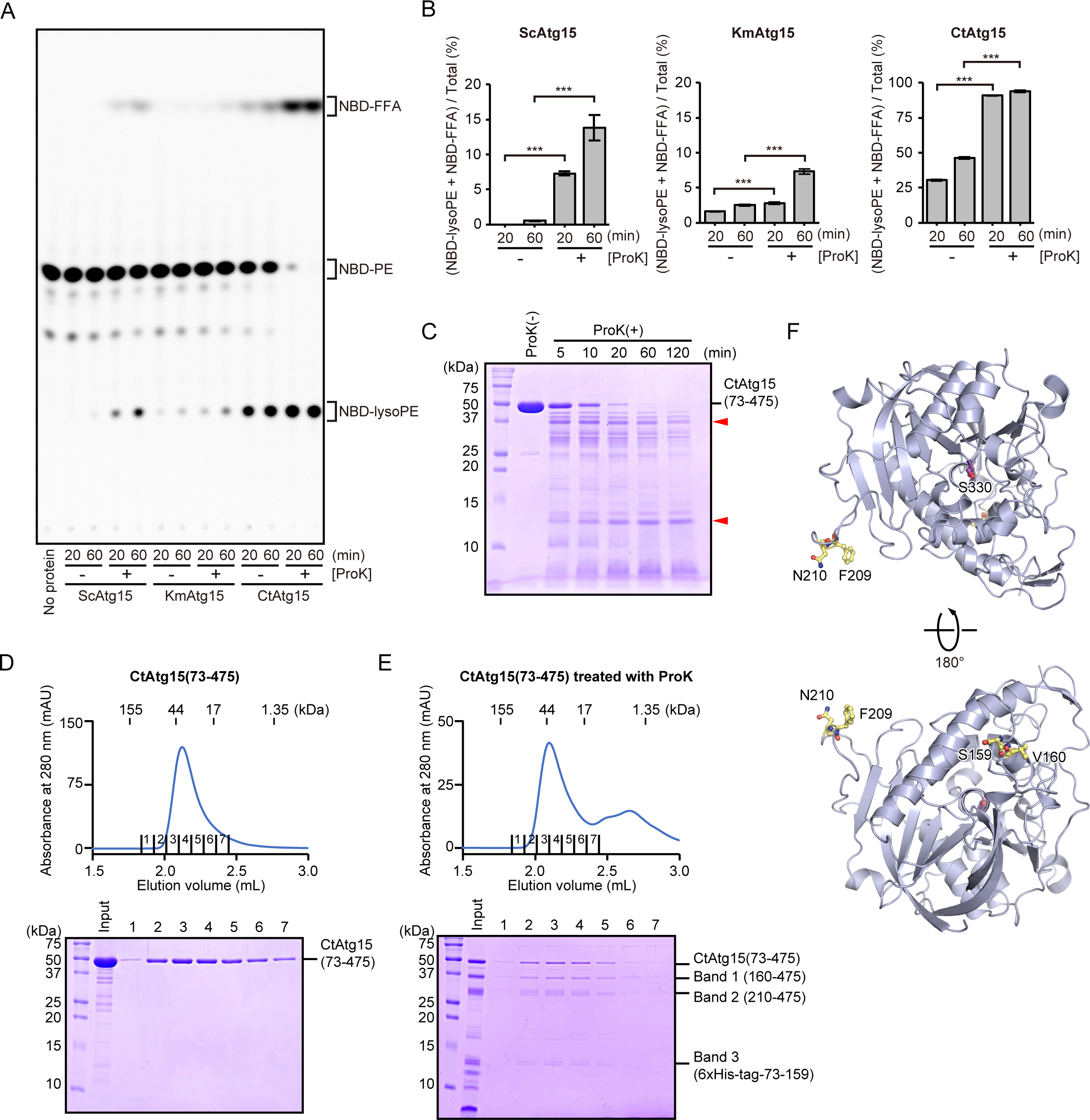
Phospholipase activity of Atg15 is increased by limited proteolysis. (A) NBD-PE and purified ScAtg15(50–471), KmAtg15(54–485) and CtAtg15(73–475) were incubated with or without proteinase K (ProK) at 30°C for the indicated times. Phospholipids were extracted and analysed by TLC. (B) Fluorescence intensities of NBD-FFA and NBD-lysoPE relative to total NBD fluorescence intensities were calculated. Values are mean ± SD (n = 3). ***: p < 0.001; p values were calculated using the unpaired two-tailed t-test. (C) Purified CtAtg15(73–475) was treated with proteinase K at 30°C for the indicated times. Proteinase K–treated and –untreated CtAtg15(73–475) were analysed by SDS-PAGE, followed by CBB staining. Stable peptides are indicated by red arrowheads. (D) Proteinase K–untreated CtAtg15(73–475) was analysed by size-exclusion chromatography using a Superdex200 increase 5/150 column (upper panel). Positions of molecular mass standards are shown above the elution profile. Fractions indicated in the elution profile were analysed by SDS-PAGE, followed by CBB staining (lower panel). (E) CtAtg15(73–475) was treated with proteinase K at 30°C for 20 min and then analysed by size-exclusion chromatography using a Superdex200 increase 5/150 column (upper panel). Positions of molecular mass standards are shown above the elution profile. Fractions indicated in the elution profile were analysed by SDS-PAGE, followed by CBB staining (lower panel). Three peptides indicated as Band 1, 2, and 3 were analysed by Edman sequencing. (F) Predicted structure of CtAtg15(73–475) by AlphaFold2. S159, V160, F209, N210, and the active-center S330 are shown using a ball-and-stick model. S159, V160, F209, and N210 are colored in yellow. S330 is colored in magenta.

### CtAtg15 is activated by cleavage between S159 and V160

Treatment of CtAtg15(73–475) with proteinase K generated two relatively stable peptides (35 kDa and 12 kDa in Fig. 2C), showing that CtAtg15(73–475) had sites that were easily cleaved by proteinase K. To investigate the conformation of the two peptides generated by limited proteolysis, we performed gel filtration analysis. This showed that the molecular mass of intact CtAtg15(73–475) was about 44 kDa, indicating that CtAtg15(73–475) is present as a monomer. After treatment with proteinase K, two peptides (bands 1 and 3) co-migrated with another peptide (band 2) to form a single peak at a similar position as that of the intact CtAtg15(73–475), suggesting that CtAtg15(73–475) retains its folding even after treatment with proteinase K (Fig. 2D–E). Edman sequencing indicated that the peptides of bands 1, 2, and 3 started at V160, N210, and the glycine residue just before the N-terminal 6×His-tag, respectively (Fig. S3). This information, together with the molecular weights shown by SDS-PAGE, demonstrated that bands 1, 2, and 3 represent 160–475, 210–475, and N-terminal 6×His-tag-fused 73–159, respectively. These results indicate that CtAtg15(73– 475) is cleaved between S159 and V160 and between F209 and N210 by treatment with proteinase K. In the AlphaFold2-predicted CtAtg15 structure, F209 and N210 are located on the exposed loop region away from the active-center S330 of CtAtg15, whereas S159 and V160 are located on opposite sides of the active center (Fig. 2F).

To investigate whether the loop region that includes F209 and N210 is important for the activity of CtAtg15, we removed the loop region from CtAtg15(73–475) to generate CtAtg15(Δ202–213), and then measured the activity. CtAtg15(Δ202–213) did not show any phospholipase activity (Fig. 3A), suggesting that the loop region that includes F209 and N210 is required for the activity of CtAtg15. Next, we examined whether cleavage between S159 and V160, or between F209 and N210, was important for the activity of CtAtg15. We inserted the human rhinovirus (HRV) 3C protease recognition sequence LEVLFQGP between S159 and V160, or between F209 and N210 (Fig. 3B), and compared CtAtg15 activity with or without treatment with the HRV 3C protease. CtAtg15 with insertion of the LEVLLFQGP sequence between F209 and N210 did not show lipase activity regardless of HRV 3C protease treatment (Fig. 3C–D), suggesting that the loop region that includes F209 and N210 needs to be intact to permit CtAtg15 activity. By contrast, the lipase activity of CtAtg15 with insertion of the LEVLFQGP sequence between S159 and V160 was higher than that of wild-type CtAtg15 without HRV 3C protease treatment (Fig. 3C–D), and this activity increased with HRV 3C protease treatment (Fig. 3C–D). These results suggest that cleavage between S159 and V160 facilitates the activity of CtAtg15.

**Figure 3.**
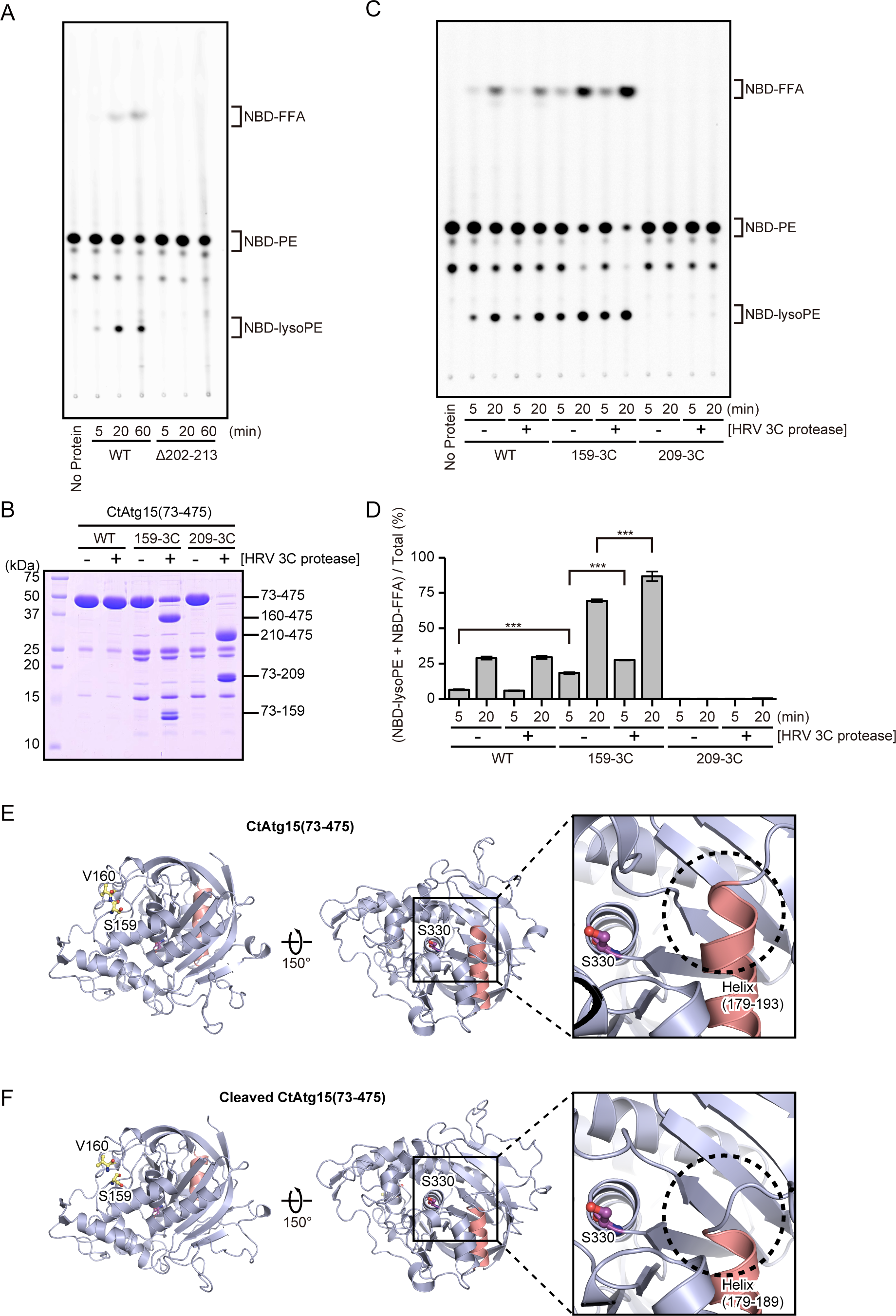
CtAtg15 is activated by the cleavage between S159 and V160. (A) NBD-PE and the purified wild-type (WT) and Δ202-213 mutant of CtAtg15(73– 475) were incubated at 30°C for the indicated times. Phospholipids were extracted and analysed by TLC. (B) WT CtAtg15(73–475) and the HRV 3C protease recognition sequence insertion mutants between S159 and V160 (159-3C) or between F209 and N210 (209-3C) of CtAtg15(73–475) were incubated with or without the HRV 3C protease at 4°C for 1h, and then analysed by SDS-PAGE, followed by CBB staining. (C) NBD-PE was incubated with the HRV 3C protease–treated or –untreated WT CtAtg15(73–475) and 159-3C or 209-3C mutants of CtAtg15(73–475) at 30°C for the indicated times. Phospholipids were extracted and analysed by TLC. (D) Fluorescence intensities of NBD-FFA and NBD-lysoPE relative to total NBD fluorescence intensities were calculated. Values are mean ± SD (n = 3). ***: p < 0.001; p values were obtained from the unpaired two-tailed t-test. (E–F) Structural change associated with the cleavage between S159 and V160. Predicted structures of (E) CtAtg15(73–475) and (F) CtAtg15(73–473) cleaved between S159 and V160. S159, V160, and the active-center S330 are shown using a ball-and-stick model. S159 and V160 are colored in yellow. S330 is colored in magenta. Magnified views around the active center of CtAtg15(73–475) are shown in the right panels. Helix regions shortened by the cleavage between S159 and V160 are colored in salmon pink and indicated with dashed circles.

We used ColabFold^23^ to predict the structural differences between CtAtg15(73– 475) and CtAtg15(73–159)+(160–475), the latter which consists of two peptides generated by cleavage between S159 and V160. Remarkably, the α-helix (179–193) near the active-center S330 in CtAtg15(73–475) became shorter after the cleavage between S159 and V160 (Fig. 3E–F). It is possible that this structural change contributes to facilitating CtAtg15 activity.

### Precursor ScAtg15 in cell lysates can be activated by limited proteolysis

We generated a ScAtg15-overexpressing plasmid as described previously^14^. Cell lysates prepared from *pep4*Δ*atg15*Δ *S. cerevisiae* cells overexpressing ScAtg15 were subjected to the lipase assay. Minimal phospholipid activity was detected in cell lysates from ScAtg15-overexpressing *pep4*Δ*atg15*Δ cells (Fig. 4A). We next examined whether ScAtg15 in *pep4*Δ cells could be activated by treatment with proteinase K. As expected, the activity of Atg15 was greatly enhanced (Fig. 4A), indicating that proteinase K treatment can compensate for the absence of Pep4. This result suggests that processing of ScAtg15 by Pep4 is important for the activation of precursor ScAtg15. By contrast, phospholipase activity was not detected in the cell lysates of ScAtg15(S332A)-overexpressing *pep4*Δ*atg15*Δ cells, even after treatment with proteinase K (Fig. 4A).

**Figure 4.**
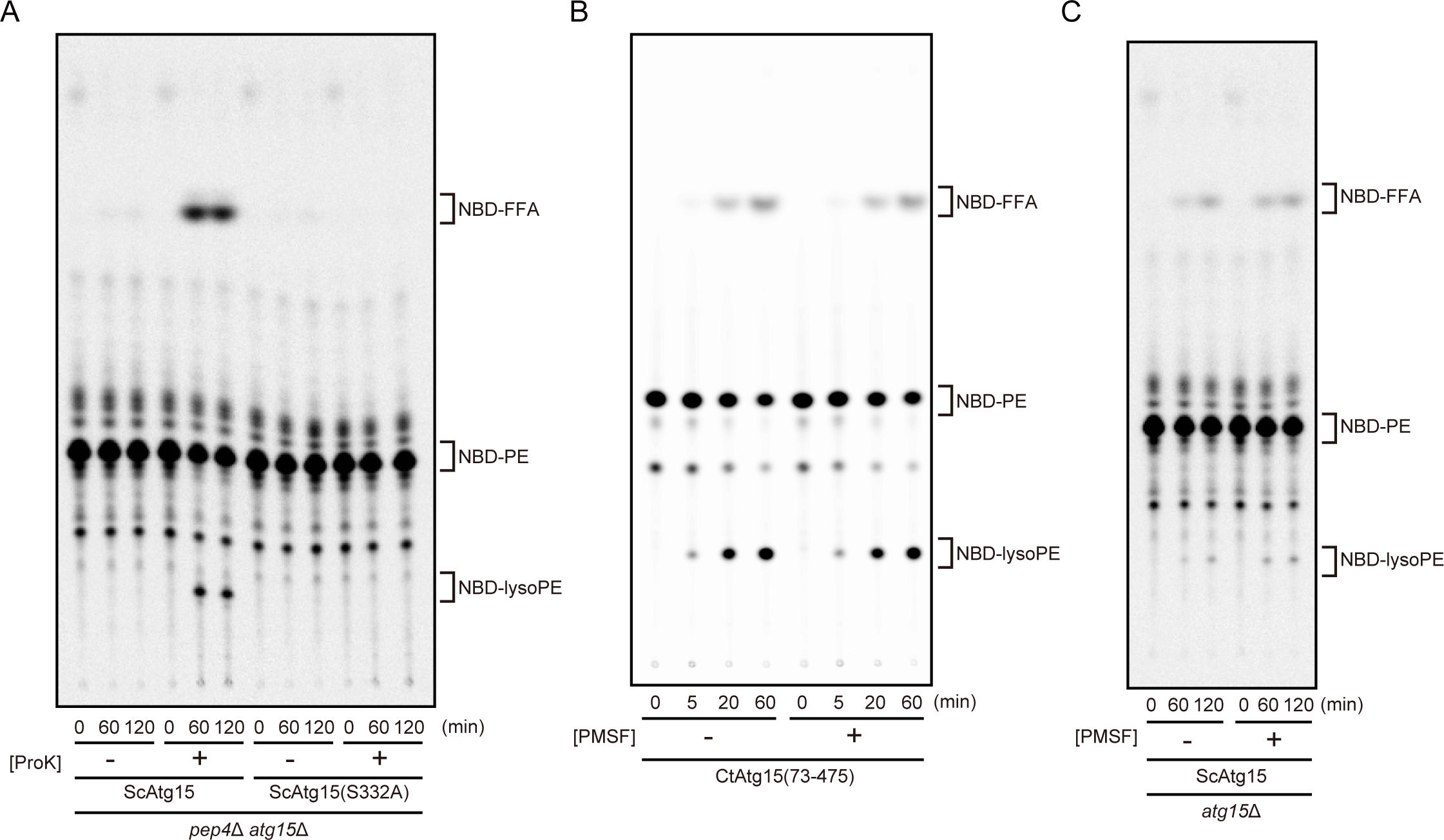
Precursor ScAtg15 in cell lysates can be activated by limited proteolysis. (A) *S. cerevisiae pep4*Δ*atg15*Δ cells harboring galactose-inducible ScAtg15 or ScAtg15(S332A) plasmids were grown in SRafCA medium to mid-log phase. ScAtg15 and ScAtg15(S332A) were overexpressed in the presence of 3% galactose for 6 h. Cell lysates were prepared by glass bead disruption. Proteinase K was added immediately before reaction onset. (B–C) Phenylmethylsulfonyl fluoride (PMSF) does not inhibit Atg15 activity. (B) NBD-PE and purified CtAtg15(73–475) were incubated with or without 1 mM PMSF at 30°C for the indicated times. Phospholipids were extracted and analysed by TLC. (C) *S. cerevisiae pep4*Δ*atg15*Δ cells harboring the galactose-inducible ScAtg15 plasmid were grown in SRafCA medium to mid-log phase. ScAtg15 was overexpressed by the addition of 3% galactose for 6 h. Cell lysates were prepared by glass bead disruption. PMSF (final 2 mM) was added every 30 min.

### Phenylmethylsulfonyl fluoride (PMSF) does not inhibit Atg15 activity

Previous studies have reported that the addition of PMSF causes the accumulation of autophagic bodies by blocking their degradation^20^. Since PMSF inhibits enzymes by irreversibly binding with a serine residue at the active center^26^, we examined whether PMSF affected the activity of Atg15. First, we demonstrated that the activity of recombinant CtAtg15 was hardly affected by the presence of PMSF (Fig. 4B). Moreover, PMSF did not seem to have any effect on the activity of Atg15 in cell lysates from *S. cerevisiae* cells (Fig. 4C). From these results, we conclude that Atg15 is not a direct target of PMSF.

### CtAtg15 can replace the activity of ScAtg15 in *S. cerevisiae* vacuoles

We next examined whether CtAtg15 compensated for the lack of *ATG15* in *S. cerevisiae* cells. To transport CtAtg15(73–475) to the vacuolar lumen, we used the GFP-fused N-terminal transmembrane domain of Pho8 (GFP-Pho8TMD), as we previously reported^16^. We fused CtAtg15(73–475) to GFP-Pho8TMD to generate GFP-Pho8TMD-CtAtg15. *S. cerevisiae atg15*Δ cells expressing GFP-Pho8TMD-ScAtg15, GFP-Pho8TMD or GFP-Pho8TMD-CtAtg15 were grown to mid-log phase and treated with rapamycin for 3 h to induce autophagy. The activity of Atg15 was estimated by the maturation of the aminopeptidase I precursor (prApe1). The propeptide in prApe1 is cleaved by vacuolar proteases; consequently, prApe1 is converted to mature Ape1 (mApe1)^8^. The decrease in the molecular weight of prApe1, as detected by immunoblot analysis, can be used as an indicator of the Atg15-mediated degradation of autophagic bodies containing prApe1. mApe1 was not detected in GFP-Pho8TMD–expressing cells, but did appear in GFP-Pho8TMD-CtAtg15–expressing cells (Fig. 5A), showing that CtAtg15 compensated for the absence of ScAtg15. This result clearly shows that CtAtg15 has the capacity to disintegrate IAC in *S. cerevisiae*.

**Figure 5.**
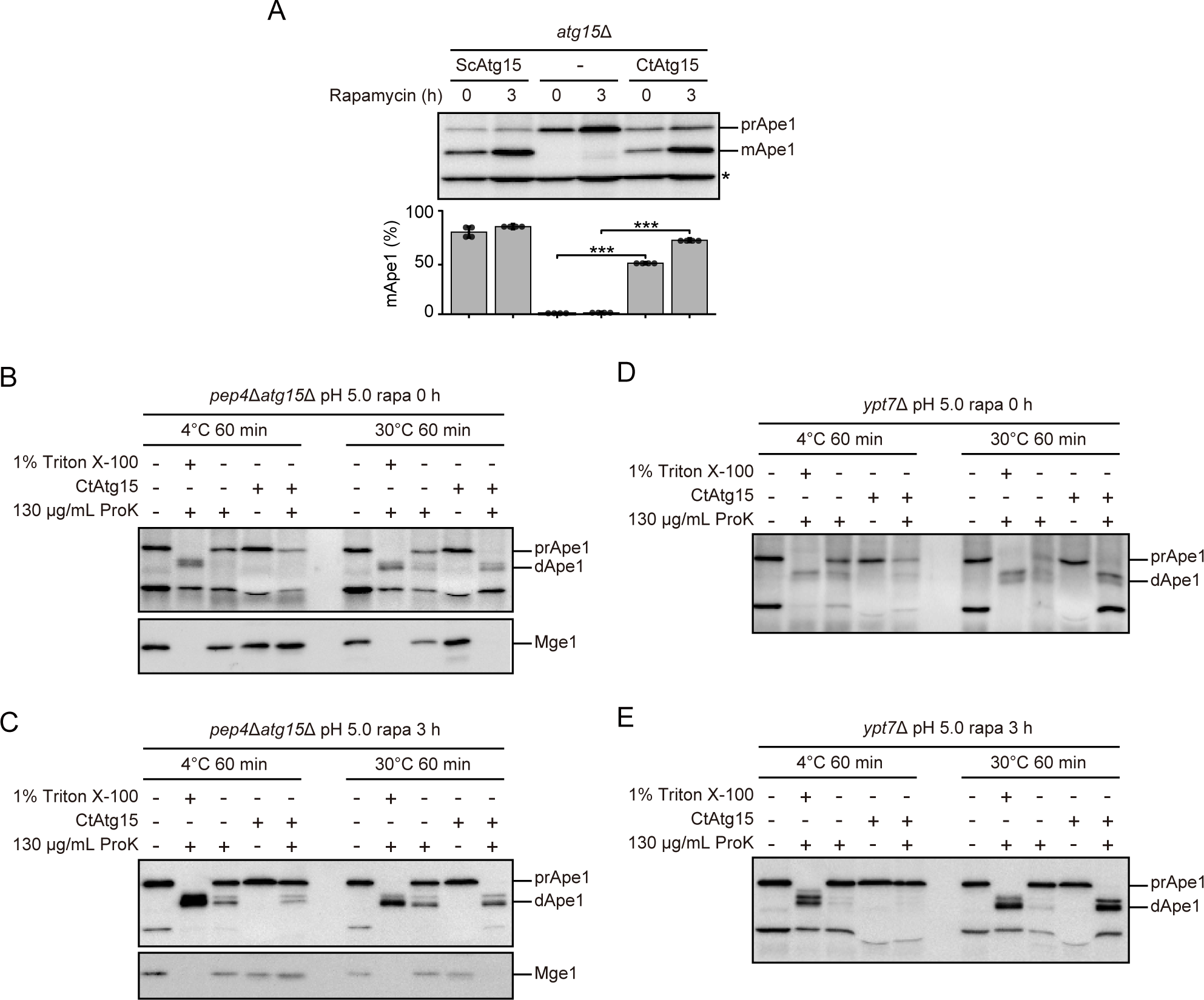
CtAtg15 disintegrates organelle membranes of *S. cerevisiae*. (A) CtAtg15 can compensate for the lack of activity of ScAtg15 in vacuoles of *S. cerevisiae*. *S. cerevisiae atg15*Δ (Y5789) cells harboring pRS316[GFP-Pho8TMD] background plasmids were grown in SDCA medium to mid-log phase, treated with rapamycin for 3 h and collected. Cell lysates equivalent to 0.15 OD_600_ units prepared by the alkaline lysis method were subjected to immunoblot analysis with anti-Ape1 antiserum. Asterisk indicates non-specific bands. Maturation of Ape1 was quantified based on the immunoblot images. ***p < 0.001 (n = 4, two-tailed Student’s t-test). (B– C) *S. cerevisiae pep4*Δ*atg15*Δ (GYS1575) cells were grown in YEPD medium to mid-log phase (B), treated with rapamycin for 3 h (C) to induce autophagy and collected. Samples were subjected to immunoblot analysis using anti-Ape1 or anti-Mge1 antisera. prApe1 and dApe1 indicate precursor Ape1 and degraded Ape1, respectively. (D–E) *S. cerevisiae ypt7*Δ (Y575) cells were grown in YEPD medium to mid-log phase (A), treated with rapamycin for 3 h (B) to induce autophagy and collected. Samples were subjected to immunoblot analysis using anti-Ape1 antiserum.

### CtAtg15 disintegrates organelle membranes in *S. cerevisiae*

To examine whether Atg15 could disintegrate organelle membranes, we performed experiments using recombinant CtAtg15 as the enzyme and the 15,000 *g* membrane fraction obtained from *S. cerevisiae* cells as the substrate. The vacuoles containing Cvt bodies were collected by centrifugation at 15,000 *g* and subjected to a proteinase K protection assay (Fig. 5B). As expected, prApe1 was resistant to proteinase K treatment at two different temperatures (Fig. 5B, lanes 3 and 8), suggesting that prApe1 is enclosed in Cvt bodies and/or vacuoles. prApe1 was degraded by simultaneous treatment with CtAtg15 and proteinase K (Fig. 5B, lane 10), suggesting that both vacuolar membranes and Cvt bodies are disintegrated by the action of CtAtg15. By contrast, prApe1 remained when the mixture of CtAtg15 and proteinase K was incubated at 4°C (Fig. 5B, lane 5), indicating that the activity of CtAtg15 was suppressed at 4°C even though proteinase K was active at that temperature (Fig. 5B, lane 2). prApe1 is selectively enclosed in autophagic bodies during autophagy. Cell lysates were prepared from *pep4*Δ*atg15*Δ cells treated with rapamycin for 3 h to induce autophagy. The vacuoles containing autophagic bodies were collected by centrifugation at 15,000 *g* and subjected to the proteinase K protection assay (Fig. 5C), yielding similar results. These findings suggest that proteinase K can access prApe1 enclosed in Cvt bodies as well as autophagic bodies inside vacuoles.

Next, we examined whether Atg15 contributes to degrading other organelle membranes. Since mitochondria are collected in the 15,000 *g* membrane fraction, we examined the degradation of mitochondrial membranes. A proteinase K protection assay using Mge1, a mitochondrial matrix protein, revealed that CtAtg15 degraded mitochondrial membranes (Fig. 5B–C). Given that Cvt vesicles do not contain mitochondria, it is likely that Atg15 can degrade mitochondrial membranes regardless of whether they are inside or outside of Cvt vesicles or autophagosomes.

Atg15 disintegrates mitochondria, which are double membrane–bound organelles. We assumed that other double membrane–bound organelles could also be degraded by Atg15. Ypt7, a Rab GTPase, is essential for vacuole fusion with Cvt vesicles or autophagosomes. In the absence of Ypt7, Cvt vesicles and autophagosomes remain localised to the cytoplasm as double membrane–bound organelles^27^. Cvt vesicles and autophagosomes were collected by centrifugation at 15,000 *g* from cell lysates of *ypt7*Δ cells and subjected to the proteinase K protection assay. The results were basically the same with those using *pep4*Δ*atg15*Δ cells (Fig. 5D–E), suggesting that Atg15 can degrade both the outer and inner membranes of Cvt vesicles and autophagosomes.

Based on the results of this study, we conclude that Atg15 can degrade not only IAC, but any organelle membrane, including mitochondria. We believe that Atg15 plays a central role in phospholipid turnover inside vacuoles through digestion of the phospholipids contained in organelle membranes.

## Discussion

### Activity of Atg15

It is known that Atg15 is transported to vacuoles via the multivesicular body pathway after synthesis^12,13,15,16^. However, it was previously unclear whether Atg15 is active during this transportation process. Previously, our group reported that the minimal essential region of ScAtg15 (residues 50–466) tagged with a transmembrane domain of Pho8 (Pho8TMD) could not disintegrate autophagic bodies in the absence of Pep4^16^. From these data, we raised two possibilities: The first was that Pho8TMD and ScAtg15(50–466) must be cleaved by Pep4, while the second was that the role of Pep4 is not restricted to the processing of Atg15, and that Pep4 cooperates with Atg15 in the disintegration of autophagic bodies^16^. In the current study we showed that both predictions were incorrect, and in fact Atg15 was activated by protease cleavage of a flexible peptide in the lipase domain (Fig. 3C). This finding strongly supports the hypothesis that Pep4, a vacuolar endopeptidase, processes the flexible peptide to activate Atg15. Thus, we think that Atg15 is inactive while being transported to the vacuole.

Vacuolar carboxypeptidase Y (Prc1) is synthesised as a precursor that is activated by Pep4-dependent cleavage^24,28,29^. Moreover, purified Prc1 can be activated by proteinase K digestion^25^. Since Atg15 was activated by proteinase K treatment (Figs. 2A–B and 4A), we assume that the precursor of Atg15 was processed by Pep4 inside vacuoles *in vivo*, similarly to Prc1.

In this study, we demonstrated that Atg15 is an enzyme that degrades organelle membranes, a so-called ‘organellase’. After synthesis, Atg15 travels to the ER, Golgi apparatus and multivesicular bodies *en route* to vacuoles^12,13,15^. It is noteworthy that Atg15 does not damage these organelle membranes, and therefore it must behave as an active phospholipase only when it reaches the vacuole. Atg15 exhibited significant activity under acidic conditions but little activity under neutral conditions (Fig. S2A). This supports our theory that Atg15 is active only inside vacuoles.

This study clearly shows that Atg15 is activated by intramolecular cleavage by proteinases to become two subunits (Fig. 2C), derived from the N-terminal region (residues 73–159) and the C-terminal region (residues 160–475), that were eluted to the same fractions by size exclusion chromatography (Fig. 2E). Previous studies have shown that full-length Atg15 is a short-lived protein with a half-life less than 1 h^12,13^. The activation of Atg15 by intramolecular cleavage might shorten its lifetime, which in turn might regulate its activity. The lifetime of Atg15 might be determined by the strength of the interaction of the two subunits and/or their stability against vacuolar proteases.

In yeast, Plb1–3 are phospholipase B enzymes predicted to localise to the plasma membrane and to the extracellular space^30,31^. Plb1 is inactive against phospholipid vesicles that form a bilayer membrane, but its activity is enhanced when the membrane is disrupted by detergents^32^, suggesting that Plb1 is unable to recognise phospholipids in membranes. In the AlphaFold2-predicted structure of the lipase domain of Plb1, the serine residue of the active center is exposed (Fig. S4A). On the other hand, in the predicted structures of both CtAtg15(73–475) and CtAtg15(73– 159)+(160–475), the serine residue of the active center is buried inside the molecule and is covered by several loop regions, including short helixes (Fig. S4B–C), suggesting that a conformational change is required for CtAtg15 activity even after intramolecular cleavage. Since CtAtg15 recognises and degrades IAC and organelle membranes (Fig. 5), the loop regions covering the serine residue of the active center might be involved in membrane recognition, and the regions might open upon membrane binding. The loop region that is cleaved between F209 and N210 by protease K is likely to be flexible and is required for CtAtg15 activity (Fig. 2F and 3A–D). The loop region containing F209 and N210 may also be involved in membrane binding of CtAtg15. The mechanism of the Atg15 conformational change upon membrane binding should be confirmed by structural analysis of Atg15 with substrate phospholipids.

### Substrate preference of Atg15

Previously, our group raised two hypotheses: first, that Atg15 may preferentially degrade specific lipids residing on the membrane of intravacuolar vesicles, and second, that it may also recognise the positive curvature of these vesicles^16^. Here we showed that Atg15 can degrade PE, PS, and PC (Fig. 1D–F), suggesting that the first hypothesis is unlikely. We performed *in vitro* experiments to determine if Atg15 could degrade actual targets, such as IAC or mitochondria, and demonstrated that the limiting membranes surrounding these organelles had disintegrated (Fig. 5B–E). This finding indicates that Atg15 disrupts organelle membranes nonspecifically, and that the substrate specificity of Atg15 is therefore weak or nonexistent.

Regarding the second hypothesis, Atg15 was able to disintegrate IAC membranes accumulated in fractionated vacuoles (Fig. 5B–C). The fact that prApe1 degradation did not occur at 4°C even in the presence of Atg15 (Fig. 5B and C, lane 5) indicates that the degradation of IAC membranes is dependent on the enzymatic activity of Atg15. Furthermore, it should be noted that the substrates used in this experiment were vacuoles containing IAC, and the fact that almost all prApe1 disappeared in the presence of Atg15 and proteinase K suggests that most vacuolar membranes are proteinase K–permeable in a manner dependent on Atg15 activity. These results indicate that externally added Atg15 can damage vacuolar membranes, and are consistent with the second hypothesis that Atg15 preferentially degrades phospholipids constituting lipid bilayers with positive curvature. However, it is also possible that intact cells are capable of repairing damaged vacuolar membranes. Currently, neither possibility can be excluded. We believe that future studies are needed to clarify whether Atg15 recognises a positive curvature. At the very least, the organelle membrane– degrading activity of Atg15 is probably regulated by multiple safety mechanisms, since Atg15 activity is highly dependent on pH and limited proteolysis dependent on a protease.

In this study, we showed that Atg15 has the capacity to disintegrate biological membranes and to digests their constituent phospholipids, thus producing lysophospholipids and free fatty acids (Fig. 1A–C). Since it is completely unknown how these degradation products are recycled and whether or not they have physiological roles, their fate should be investigated in future studies. We believe that the physiological significance of organelle membrane degradation revealed by the study of Atg15 will be useful to clarify the importance of this degradation in other eukaryotic cells.

## Materials and Methods

### Strains and media

Shuffle T7 Express Competent *E. coli* cells (New England Biolabs) were used for expression of recombinant Atg15.

The yeast strains used in this study are listed in Table 1. The *PEP4* gene was disrupted by DNA fragments amplified by polymerase chain reaction (PCR) from plasmid pFA6a-*hphNT1*^33^. Yeast cells were cultured in YEPD medium (1% Bacto yeast extract, 2% Bacto peptone, and 2% glucose) and SDCA medium (0.17% Difco yeast nitrogen base without amino acids and ammonium sulfate, 0.5% ammonium sulfate, 0.5% Bacto casamino acids, and 2% glucose), or SRafCA (0.17% Difco yeast nitrogen base without amino acids and ammonium sulfate, 0.5% ammonium sulfate, 0.5% Bacto casamino acids, and 2% raffinose) supplemented with appropriate nutrients for plasmid selection. Reagents were purchased from Fujifilm Wako Pure Chemical unless otherwise noted.

**Table 1.** Yeast strains used in this study.

| Name | Genotype | Source | Figures |
| --- | --- | --- | --- |
| BY4741 | *MATa leu2Δ0 ura3Δ0 his3Δ1 met15Δ0* | Ref 34 |  |
| Y5789 | BY4741; *atg15Δ::kanMX* | Open Biosystems | 4C, 5A |
| GYS1575 | BY4741; *atg15Δ::kanMX pep4Δ::hphNT1* | This study | 4A, 5B, 5C |
| Y575 | BY4741; *ypt7Δ::kanMX* | Open Biosystems | 5D, 5E |

### Plasmids

The plasmids used in this study are listed in Table 2. To construct *E. coli* expression plasmids for ScAtg15(50–471) and KmAtg15(54–485) with the N-terminal 6×His tag, the DNA fragments were amplified by PCR from the genomes of *S. cerevisiae* and *K. marxianus*, respectively, and cloned into a pETDuet-1 vector using the *Bam*HI and *Xho*I sites. To construct an *E. coli* expression plasmid for CtAtg15(73– 475) with the N-terminal 6×His tag, the synthetic gene optimised for *E. coli* codon usage was obtained from Eurofins Genomics and cloned into a pETDuet-1 vector using the *Bam*HI and *Xho*I sites. The expression plasmids for CtAtg15(Δ202–213) and CtAtg15(73–475) were constructed by inverse PCR and inserted into the HRV 3C protease recognition sequence (LEVLFQGP) between S159 and V160, or between F209 and N210. For construction of the Atg15 overexpression plasmid for yeast, the PCR-amplified *ATG15* gene was cloned into the pYES2/NT C vector (Thermo Fisher Scientific) using the *Bam*HI and *Xho*I sites as described previously^14^. Point mutants were generated by site-directed mutagenesis. For the complementation test of CtAtg15 in yeast, CtAtg15(73–475) was expressed from a pRS316-YEGFP-Pho8TMD-Atg15C background plasmid^16^ by substituting the CtAtg15(73–475) region for the ScAtg15(50–520) region using the NEBuilder HiFi DNA Assembly Cloning Kit (New England Biolabs). All of the constructs were sequenced to confirm their identities.

**Table 2.**
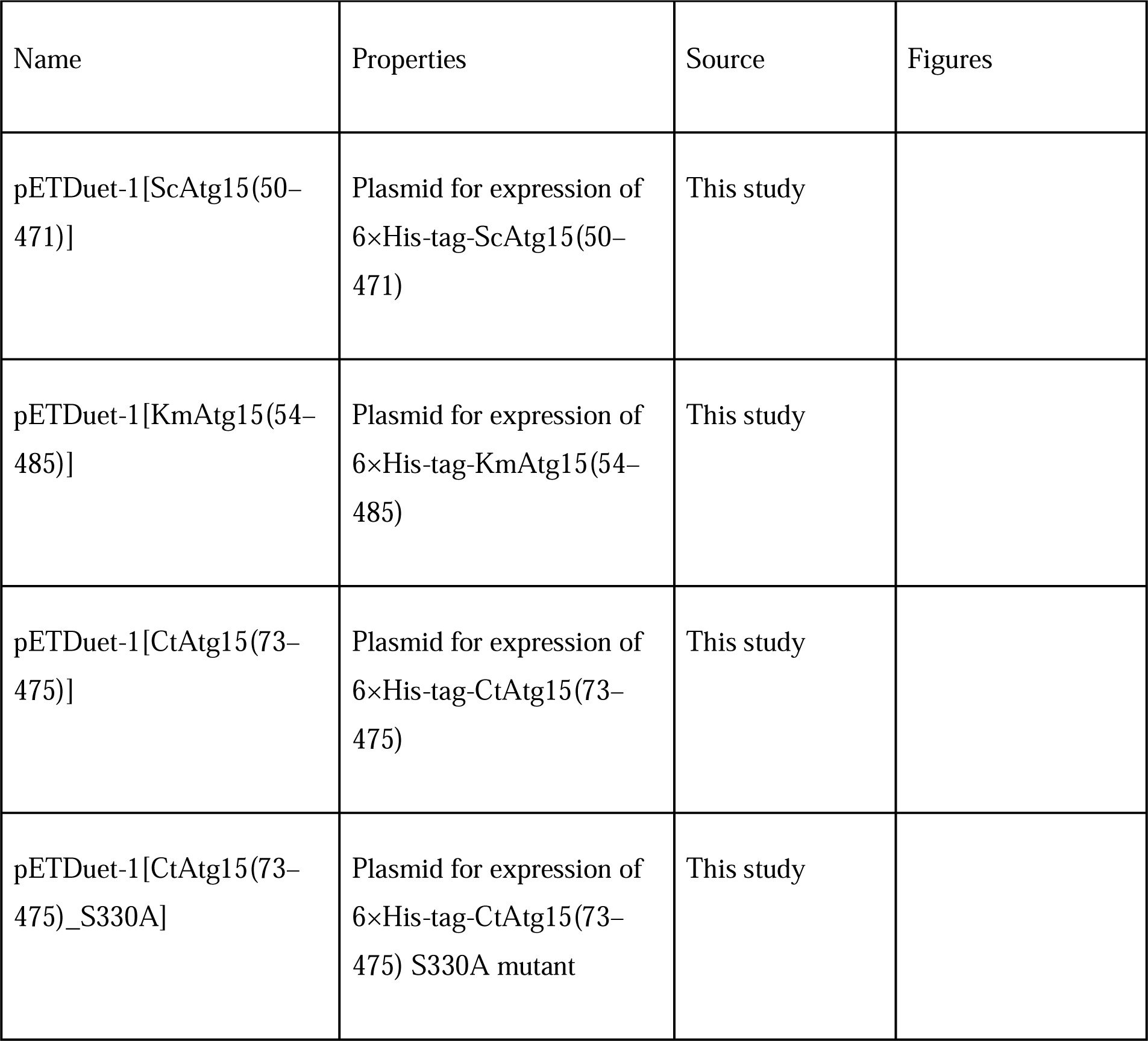
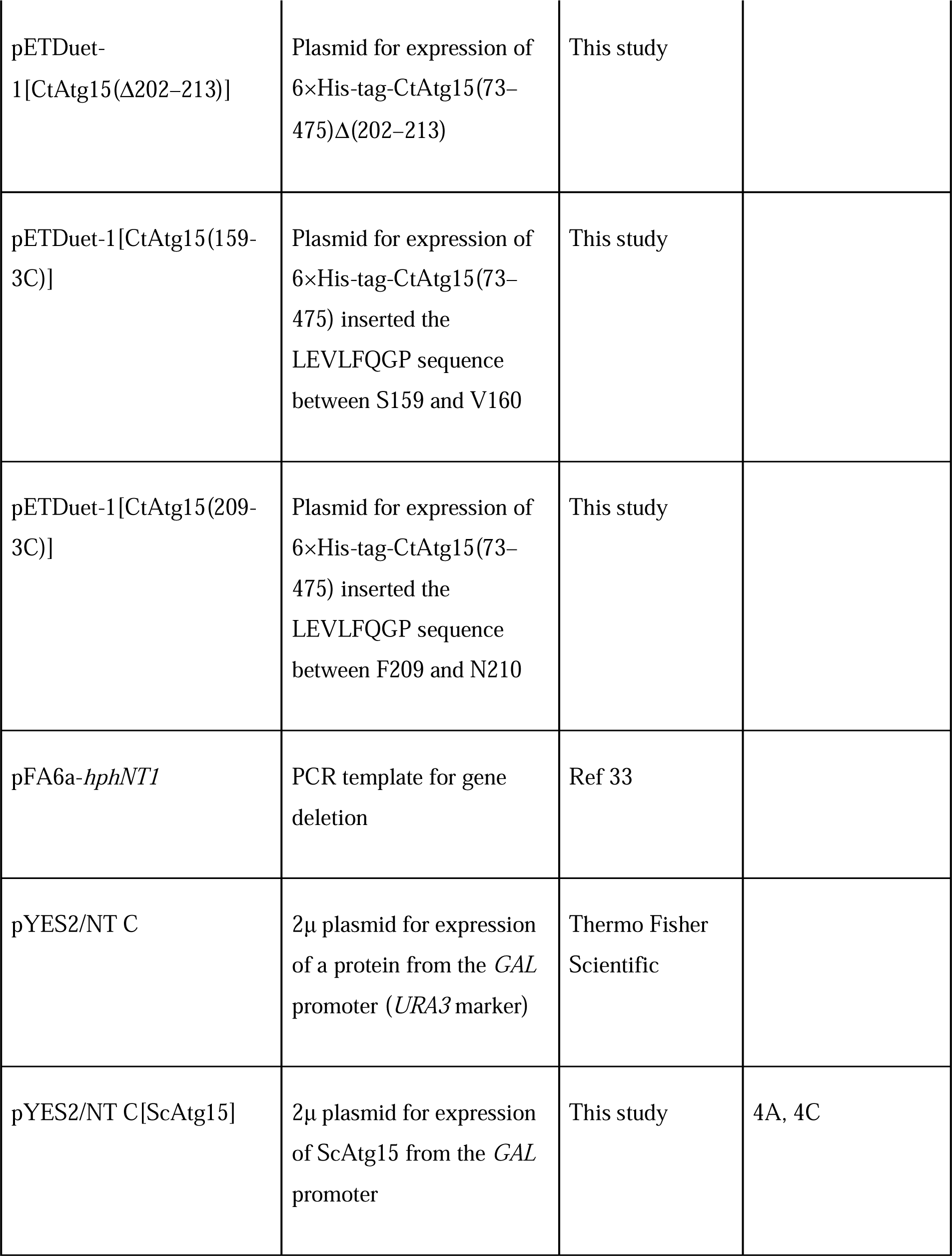
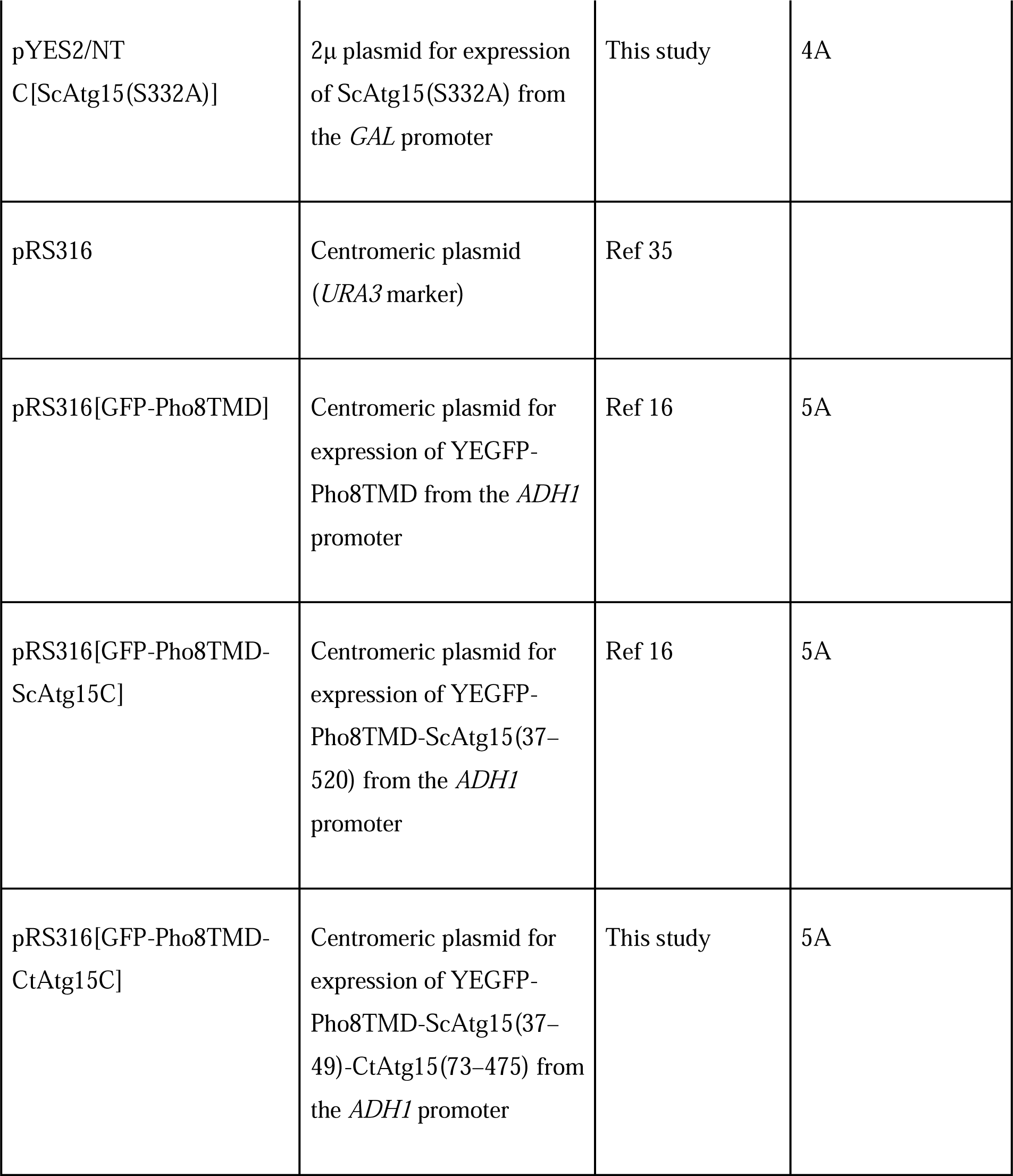
Plasmids used in this study.

### Purification of recombinant Atg15

Target proteins were expressed in *E. coli* SHuffle T7 cells cultured in LB medium supplemented with 100 μg/mL ampicillin at 37°C. When the optical density of the culture at 600 nm reached ∼0.8, isopropyl-β-D-thiogalactopyranoside (IPTG) was added to a final concentration of 0.1 mM, and cultured at 16°C for 20 h to induce protein expression. After induction of protein expression, the cells were harvested, resuspended in lysis buffer (50 mM Tris-HCl [pH 8.0], 500 mM NaCl, and 20 mM imidazole), disrupted by sonication and centrifuged to pellet the insoluble debris. The supernatant was loaded onto a Ni-NTA Agarose column (QIAGEN) equilibrated with lysis buffer. The column was washed with lysis buffer, and His-tagged proteins were eluted using elution buffer (50 mM Tris-HCl [pH 8.0], 100 mM NaCl, and 250 mM imidazole). Proteins were further purified by size exclusion chromatography using a Superdex 200 10/300 increase column (Cytiva), with an elution buffer of 20 mM Tris-HCl (pH 8.0) and 150 mM NaCl.

### Phospholipids and liposomes

The phospholipids 18:1-12:0 NBD-PE (1-oleoyl-2-{12-[(7-nitro-2-1,3-benzoxadiazol-4-yl)amino]dodecanoyl}-sn-glycero-3-phosphoethanolamine, 810156C), 18:1-12:0 NBD-PS (1-oleoyl-2-{12-[(7-nitro-2-1,3-benzoxadiazol-4-yl)amino]dodecanoyl}-sn-glycero-3-phosphoserine, 810195C), 18:1-12:0 NBD-PC (1-oleoyl-2-[12-[(7-nitro-2-1,3-benzoxadiazol-4-yl)amino]dodecanoyl]-sn-glycero-3-phosphocholine, 810133C), DOPC (1,2-dioleoyl-sn-glycero-3-phosphocholine, 850375C), and DOPE (1,2-dioleoyl-sn-glycero-3-phosphoethanolamine, 850725C) were obtained from Avanti Polar Lipids, Inc. Lipids in stock solutions in chloroform were mixed at a molar ratio of DOPC/DOPE/18:1-12:0 NBD-PE = 50/40/10, and the solvent was evaporated. The lipid film was hydrated in 50 mM acetate buffer (pH 4.0) supplemented with 150 mM NaCl. The lipid suspension was incubated at room temperature for 30 min and extruded 20 times through a polycarbonate 0.4-μm filter using a mini-extruder (Avanti Polar Lipids, Inc.).

### Thin-layer chromatography (TLC)

Eight micromolar NBD-labeled phospholipids were incubated with 20 μM purified ScAtg15(50–471), KmAtg15(54–485) and CtAtg15(73–475) or its mutants in 50 μL assay buffer (50 mM acetate buffer [pH 4.0], 150 mM NaCl, and 0.2% Triton X-100) at 30°C. To measure the activities of CtAtg15(73–475) at various pH conditions, the reactions were carried out in different buffers as follows: 50 mM acetate buffer (pH 3.1, 4.0, and 5.0) and 50 mM MES buffer (pH 6.0). These buffers were supplemented with 150 mM NaCl and 0.2% Triton X-100. Eight micromolar NBD-PE was incubated with 20 μM purified CtAtg15(73–475) in 50-μL volumes of different buffers at 30°C. To measure the activity of CtAtg15(73–475) against NBD-PE in liposomes, NBD-PE– containing liposomes (80 μM; DOPC/DOPE/18:1-12:0 NBD-PE = 50/40/10) were incubated with 20 μM purified CtAtg15(73–475) in 50 μL assay buffer without Triton X-100 at 30°C. To measure the activities of Atg15 with limited proteolysis, 8 μM NBD-PE was incubated with 20 μM purified Atg15 proteins and 6.5 μg proteinase K in 50 μL assay buffer at 30°C. To measure the activities of the HRV 3C protease–treated CtAtg15(73–473) mutant, 100 μM CtAtg15(73–475) mutant was treated with 1.5 μM HRV 3C protease at 4°C for 1 h before incubation with NBD-PE. After incubation for the indicated time periods, lipid extraction was performed using the Bligh and Dyer method with modifications. Briefly, 750 μL of 2:1 chloroform/methanol was added to the samples and vortexed for 10 min to extract lipids. One hundred microliters of water was added to the samples, which were then vortexed for 10 min. The organic phase was separated by centrifugation at 1,000 *g* for 2 min, collected, and dried under N_2_ gas. The resulting lipid film was dissolved in chloroform and analysed by TLC using chloroform/ethanol/water (65:25:4) as a developing solvent. NBD signals were detected and quantified using a ChemiDoc imaging system with ImageLab software (Bio-Rad). TLC plates were purchased from Merck Millipore.

Yeast cells harboring galactose-inducible ScAtg15 plasmids were grown to approximately 6 × 10^6^ cells/mL in SRafCA(-U) medium. After addition of galactose at the final concentration of 3%, cells were grown overnight. Cells were collected and resuspended in a reaction buffer (100 mM MES [pH 6.0]) and lysed using a Beads shocker (Yasui Kikai; speed 2,500 rpm, total time 720 s, on time 10 s, off time 30 s) in 2 mL tubes. Lysates were cleared at 380 *g* for 1 min at 4°C. Supernatants were used for reactions as described above.

### Mass spectrometry

After NBD-PE and products produced by CtAtg15(73–475) were separated by TLC, they were scraped from the TLC plate. The scraped samples were suspended in 50 μL of water. The suspended samples were mixed with 750 μL of 2:1 chloroform/methanol and then vortexed for 10 min to extract lipids. One hundred microliters of water was added to the samples, which were then vortexed for 10 min. The organic phase was separated by centrifugation at 1,000 *g* for 2 min, collected, and dried under N_2_ gas. The resulting lipid film was dissolved in 0.5 mL of methanol containing 10 mM ammonium acetate. Lipid samples were injected into a Turbo V electrospray ion source using a syringe pump with a flow rate of 10 μL/min, and analysed by direct infusion mass spectrometry using a 3200 QTrap system (SCIEX). Lipids were ionized using negative ion mode electrospray ionisation, and ionised fragments were detected by mass spectrometry.

### Limited proteolysis

Ninety micromolar CtAtg15(73–475) was treated with 2.6 μg proteinase K in 50 mM acetate buffer (pH 4.0) and 150 mM NaCl at 30°C for 20 min. After incubation, the proteolysis reaction was stopped by the addition of 4-(2-aminoethyl)benzenesulfonyl fluoride at a final concentration of 1 mM. The CtAtg15(73–475) before and after limited proteolysis was subjected to size exclusion chromatography using a Superdex 200 5/150 increase column (Cytiva), with an elution buffer of 20 mM Tris-HCl (pH 8.0) and 150 mM NaCl. Each fraction analysed by size exclusion chromatography was subjected to SDS-PAGE, followed by CBB staining. Proteins separated by SDS-PAGE were transferred to Immobilon-P membranes (Merck Millipore). Three major bands on the membrane were cut and subjected to N-terminal Edman sequencing by Hokkaido System Science Co., Ltd.

### Proteinase K protection assay

Yeast cells were grown to log phase, collected, converted to spheroplasts and mechanically disrupted with 3.0-μm polycarbonate filters (Whatman) in lysis buffer (HES_1.0_ buffer: 20 mM HEPES [pH 7.5], 5 mM EDTA, and 1 M sorbitol). After cell debris was removed by centrifugation, cell lysates were centrifuged at 15,000 *g* for 15 min. The pellets were resuspended in AES_1.0_ buffer (80 mM potassium acetate [pH 5.0], 5 mM EDTA and 1 M sorbitol), divided into aliquots and treated with or without 20 μM CtAtg15(73-475), 1% Triton X-100, and proteinase K (9034, Takara Bio). The reactions were terminated by addition of 15% trichloroacetic acid. Precipitants were washed with acetone and dissolved in SDS-PAGE sample buffer containing 4 mM Pefabloc SC (11429876001, Roche). Equivalent protein amounts were subjected to immunoblotting analysis.

### Immunoblotting analysis

Proteins separated by SDS-PAGE were transferred to Immobilon-P membranes (Merck Millipore) utilizing a Trans-Blot SD Semi-Dry Transfer Cell (Bio-Rad). Following transfer, membranes were blocked with 2% skim milk in Tris-buffered saline containing 0.05% Tween 20 (Nacalai Tesque) for 30 min at room temperature. Membranes were then incubated with anti-GFP antibody (JL-8; Clontech) at a dilution of 1:5,000, or with anti-Ape1 antiserum at a dilution of 1:10,000^5^ for 60 min at room temperature. Subsequently, the membranes were washed once and treated with horseradish peroxidase–labeled anti-rabbit or anti-mouse secondary antibodies (Jackson ImmunoResearch) at a dilution of 1:5,000 for 30 min, followed by three washes. Chemiluminescent signals generated using ECL Prime (Cytiva) or ImmunoStar LD were detected on an IR-LAS 1000 imaging system.

## Abbreviations

CBB: Coomassie brilliant blue
FFA: Free fatty acid
HRV: Human rhinovirus
IAC: Intervacuolar autophagic compartments
NBD: Nitrobenzoxadiazole
PC: Phosphatidylcholine
PE: Phosphatidylethanolamine
PMSF: Phenylmethylsulfonyl fluoride
SDCA: Synthetic dextrose plus casamino acid
SDS-PAGE: Sodium dodecyl sulfate–polyacrylamide gel electrophoresis
SRafCA: Synthetic raffinose plus casamino acid
TLC: Thin-layer chromatography
TMD: Transmembrane domain

## Acknowledgements

We thank Dr Yasushi Tamura for his valuable advice and for use of the equipment involved in protein preparation. We also thank Drs Toshiya Endo and Koji Okamoto for Mge1 antiserum. This work was supported by a grant from the Naito Foundation (to K. Suzuki) and by Grants-in-Aid for Scientific Research from the Ministry of Education, Culture, Sports, Science and Technology of Japan (22K06096 and 22H02569 to YW; 20H05313, 21K19205, and 22H02569 to K. Suzuki). This work was supported by CREST, the Japan Science and Technology Agency, Japan (JP201032912 to K. Suzuki).

## Author contributions

YW and K. Suzuki, conceptualization; YW and K. Suzuki, methodology; YW and K. Suzuki, validation; YW, YI, K. Sasaki, and K. Suzuki, formal analysis; YW, YI, K. Sasaki, and K. Suzuki, investigation; YW, YI, and K. Suzuki, resources; YW and K. Suzuki, writing–original draft; YW and K. Suzuki, writing–review and editing; YW and K. Suzuki, project administration management and coordination responsibility; YW and K. Suzuki, funding acquisition.

---

**Figure S1. Predicted structures of Atg15 homologs.**

Predicted structures of (A) *S. cerevisiae* Atg15 (ScAtg15), (B) *K. marxianus* Atg15 (KmAtg15), and (C) *C. thermophilum* Atg15 (CtAtg15) are shown in ribbon diagrams. Putative lipase domains of ScAtg15(50–471), KmAtg15(54–485), and CtAtg15(73– 475) are colored in light blue. The remaining unstructured N- and C-terminal regions are colored in yellow. (C) Sequence alignment between ScAtg15, KmAtg15, and CtAtg15 by ClustalW. The conserved residues and type-conserved residues are marked by asterisks and dots, respectively. The serine residues 332, 336, and 330 of ScAtg15, KmAtg15, and CtAtg15, respectively, that were predicted as an active center are shaded in black.

**Figure S2. Characterisation of CtAtg15.**

(A) pH dependence of CtAtg15. NBD-PE and purified CtAtg15(73–475) were incubated at 30°C under different pH conditions for the indicated times. Phospholipids were extracted and analysed by TLC. (B) The activity of CtAtg15 against NBD-PE in liposomes. NBD-PE in Triton X-100 micelles or NBD-PE-containing liposomes (DOPC/DOPE/18:1-12:0 NBD-PE = 50/40/10) were incubated with the wild-type (WT) or S330A mutant of CtAtg15(73–475) at 30°C for the indicated times. Phospholipids were extracted and analysed by TLC.

**Figure S3. Cleavage sites by protease K determined by Edman sequencing.**

Amino acid sequence of CtAtg15(73–475) with 6×His-tag fused to the N-terminus. The N-terminal residues of the three peptides indicated as Bands 1, 2, and 3 in Fig. 2E are indicated by red arrowheads.

**Figure S4. Predicted structures of ScPlb1 and CtAtg15.**

Predicted structures of (A) the lipase domain of *S. cerevisiae* Plb1(35–586), (B) CtAtg15(73–475), and (C) CtAtg15(73–473) cleaved between S159 and V160 are shown in ribbon diagrams (left) and in surface representations (right). The active-center serine residues, S147 of ScPlb1 and S330 of CtAtg15, are shown using a ball-and-stick model and colored in magenta. The loop regions near the active center are colored in yellow.

