## Supplementary material for "Atg15 is a vacuolar phospholipase that disintegrates organelle membranes": Fig. S1

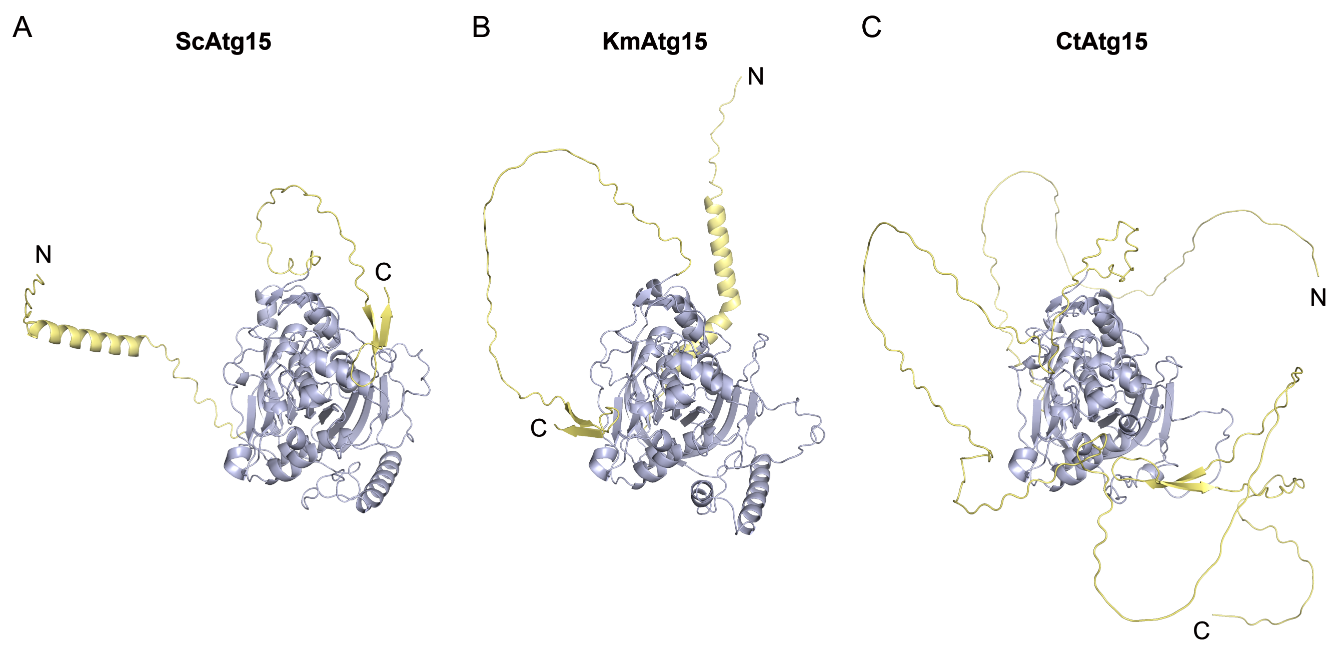


D

```
ScAtg15       1 MLHKSPSRKRFASP----LHLGCILTLTVLCLIAYYFALPDYLSVGKSSSRGAMDQKSDG 56
KmAtg15       1 MKPKTTPRKRYSAKNASLVTLLIVLTLYLYLNKDIIESKYHRNDKFKDTHAVTADTTNLH 60
CtAtg15       1 MVATSSEGGATTTCASARRVTAHRLWTSVLALSAPFLPAAAVAAGNSNGRADAQAVLPID 60
                *   :      ::           *   :                       :

ScAtg15      57 TFRLKSIYRHGVGAN-HRLHQRLEVTPEVISAAGMLYQETTTQG----QDFEDQEPLWTT 111
KmAtg15      61 TLRIKEIHFRPFNSTGRRKYGQVILTPELVADAKDRFSEAVSVSGSRSEGQEEDDAYDTK 120
CtAtg15      61 APAIPPLMPEPPAPAEHVFTLRHIFHHGTYKHPGLHRKLDIPKSYAQTLWLKAEDGYDAM 120
                :  :  :         :    :                             : ::   :

ScAtg15     112 NAEYATTNPFDFEFELRRMPLLMKRMKERDPEFIESYIYG--ETYMTEEEEHAMWIDDDI 169
KmAtg15     121 LWSKSDSNPWTHEFKLKQQTIQMTRMAYRDPDYVESFLDFANENPTMAQKVHLEWVDENV 180
CtAtg15     121 PIG---------PLRARSNSVQIQRLADRRPSVVDPMVAEARQRGYVSVMSPSAWRLDTV 171
                             :  :   : : *:  * *  ::  :    :           *  : :

ScAtg15     170 VAPNITDRGTVVSLALMSSNAYVRIPQTGDWRNVT---EPWNETEPEDFGWDGDGIRGHV 226
KmAtg15     181 LAPNVTDKDTIISLALMSSNAYVPLPFEGDWRNISGWDHDANPDFPDGIGWDADGVRGHI 240
CtAtg15     172 PGPNVTDKETILSLAYMSADAYVQTPNEADWEDVG-----SGFNRSVDFGWQSDGLRGHV 226
                  **:**: *::*** **::***  *   ** ::              :**: **:***:

ScAtg15     227 FYNEVENIVVLSIKGTSAQGLPGSGEDETTGNDKINDNLLFSCCCARVS-YLWTTVCDCY 285
KmAtg15     241 FTNEDSSVVVIAIKGTSAQGLPGSGDDATTENDKLNDNLLFSCCCARVS-YLWTTACDCY 299
CtAtg15     227 FADETNSTIVIGLKGTSLAVFDGDG---TSTNDKVNDNLFFSCCCAQQGPWTWRQVCDCA 283
                * :*    :*: :****   : * *   *: ***:****:******:   : *   ***

                                                              ▼
ScAtg15     286 VKSYICDESCLEKELRRKDRFYSAVVDIYKGVLKEYPDAAIWVTGHSLGGALASLLGRTF 345
KmAtg15     300 VKSYTCDETCLEKELNRKDRYYQAVLDIYRTVIEDHPNAAIWITGHSLGGALASLLGRTF 359
CtAtg15     284 TGTYTCNNTCVREALRQENRYYQAARELYSNVTELYPNSNVWITGHSLGGAVSSFLGLTY 343
                  :* *:::*: : * :::*:*  ::*  * : :*:: :*:********::*:** *:

ScAtg15     346 GLPAVAFESPGELLPSKRLHLPFPPGLPS------YMEGIWHFGHNADPIFMGTCNGASS 399
KmAtg15     360 GAPAVAFEAPGELLATRRLHLPIPPGLPS------YQEGIWHIGHTADPIFMGTCNGASS 413
CtAtg15     344 GVPVVTFQAVPEALPASRLGLPVPPGADPNHPQMRNYTGAFHFGHTADPVYMGTCNGATA 403
                * * *:*::  * * : ** ** ***            * :*:** ***::*******::

ScAtg15     400 SCSLVGYAMETACHTGRVCVYDVVNDKGWSVNMFNHRIHKVIDEVLLGYEQAAKCVEPEP 459
KmAtg15     414 SCSIAGYAMETSCHSGKVCIYDVVTDKGWHVNMLNHRIHTVIDGVLNDYENVAKCEVPEP 473
CtAtg15     404 SCSFGGYAMETTCHTGMECVYDTVGDKGWRVGIGTHKIRAVIQDVIMKYDTVPECKPTPE 463
                ***: ******:**:*  *:** * **** * :  *:*: **: *:  *:   :*

ScAtg15     460 CVDCYNWKFIPSRDWESSSRLITKTKSHAA---------------------------PTT 492
KmAtg15     474 CNDCFNWKYIRGRDEPHSSSVTSSSSATNK---------------------------KPT 506
CtAtg15     464 CRDCGSWKMYESNGTETTTRSSSTSMATKTRTRTETCKTPGWWGCLDETTTTGEVTKPPT 523
                * **  **         ::   : : :                                *

ScAtg15     493 TTRTTATT---------------------------------------------------- 500
KmAtg15     507 TSVFTSTTSLTATTSSA------------------------------------------- 523
CtAtg15     524 TTTTTTTTTTTTTTTTTTCHTPGWFGCKDKTTTETTTSTTCLTPGRFWGCYDETTTTAAS 583
                *:  *:**

ScAtg15     501 -------------------------------TSSSTCVGRNWLGFCTKYEL--------- 520
KmAtg15     524 ------------------------------EKTTSSCIGRNWIGICTRYGI--------- 524
CtAtg15     584 TITTTRPLDSTITPPPAVPTTTSTPTTTEEPPASSTCVERTWYGVCKEWEEVDPTPTKVP 643
                                                ::*:*: * * * *  :

ScAtg15     520 ------------------------------ 520
KmAtg15     524 ------------------------------ 524
CtAtg15     644 EPSQTCVERTWWGWCREWEGGDEMVRKEDM 673
```

Figure S1
