## Supplementary figures and images for "Atg15 is a vacuolar phospholipase that disintegrates organelle membranes"

### Fig. S2

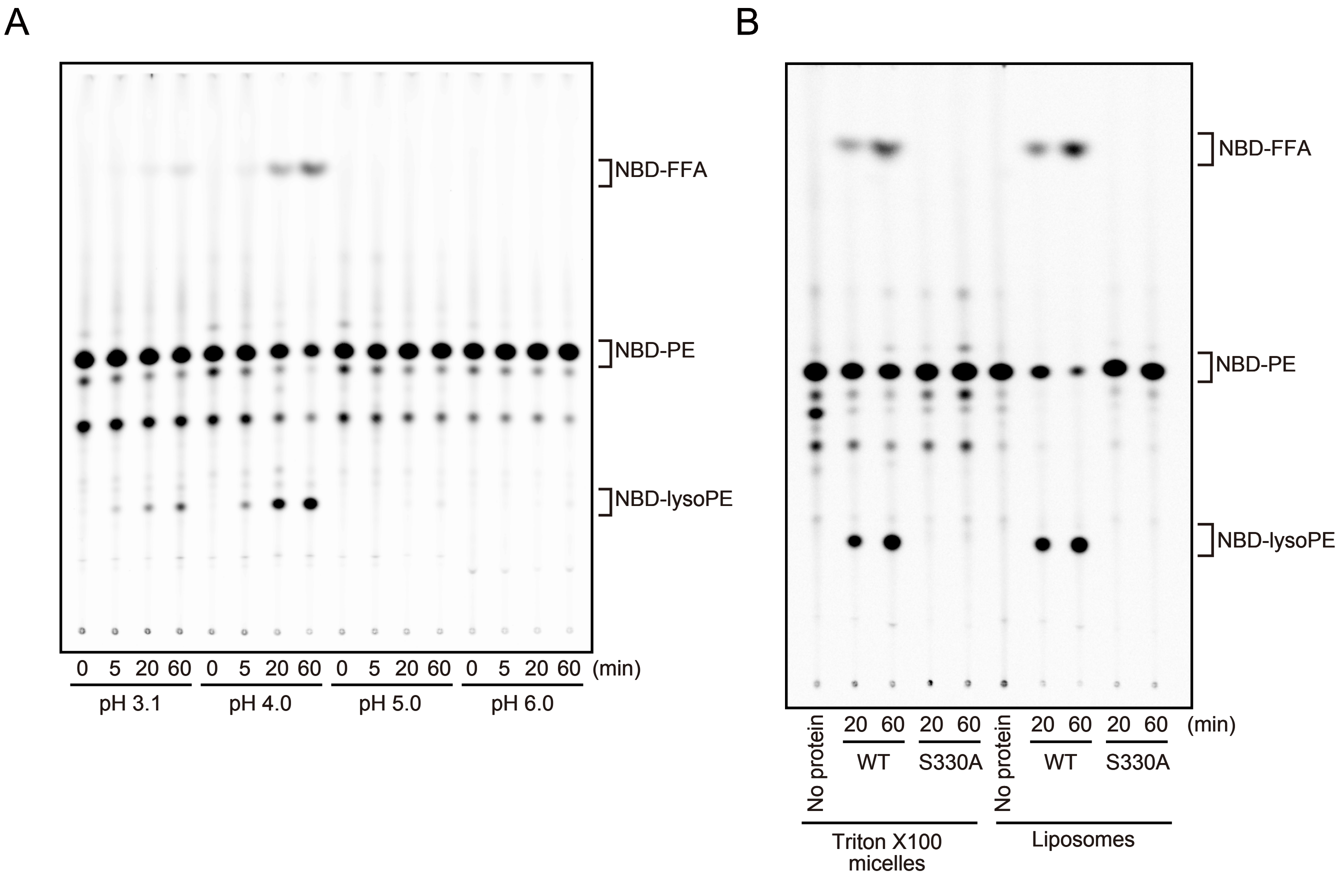

### Fig. S4

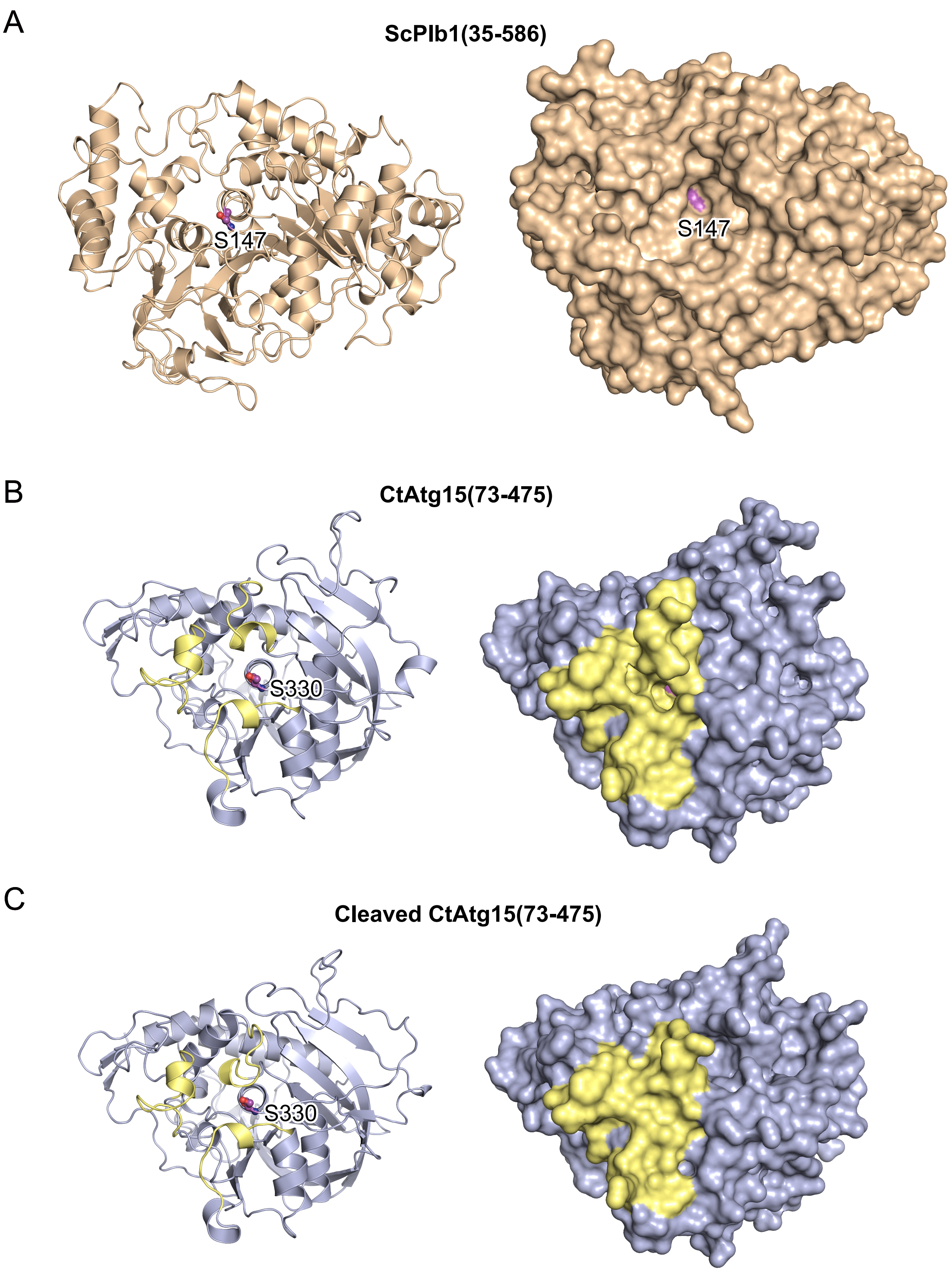
