## Supplementary material for "Atg15 is a vacuolar phospholipase that disintegrates organelle membranes": Fig. S3

Band 3

MGSSHHHHHHSQDP

73 APAEHVFTLRHIFHHGTYKHPGLHRKLDIPKSYAQTLWLKAEDGYDAMPIGPLRARSNSVQIQRLADRRPSVVDPMVAEA 152

Band 1 Band 2

153 RQRGYVSVMSPSAWRLDTVPGPNVTDKETILSLAYMSADAYVQTPNEADWEDVGSGFNRSVDFGWQSDGLRGHVFADETN 232

233 STIVIGLKGTSLAVFDGDGTSTNDKVNDNLFFSCCCAQQGPWTWRQVCDCATGTYTCNNTCVREALRQENRYYQAARELY 312

313 SNVTELYPNSNVWITGHSLGGAVSSFLGLTYGVPVVTFQAVPEALPASRLGLPVPPGADPNHPQMRNYTGAFHFGHTADP 392

393 VYMGTCNGATASCSFGGYAMETTCHTGMECVYDTVGDKGWRVGIGTHKIRAVIQDVIMKYDTVPECKPTPECRDCGSWKM 472

473 YES 475

Figure S3
